# RIPK3-dependent necroptosis is the primary cause of RIPK1-deficient-induced immunodeficiency

**DOI:** 10.1101/2023.02.22.526137

**Authors:** Xiaoyan Liu, Xinxin Zhu, Zhaoqian Shu, Liming Sun, Huayi Wang

## Abstract

RIPK1 (receptor-interacting serine/threonine kinase 1) plays a pivotal role in developing the immune system—patients with homozygous loss-of-function mutations of RIPK1 present with immunodeficiency and intestinal inflammation. Here, we reported that RIPK1-deficient rats represented defects of human patients and were certified as a suitable disease model. Knocking out *Ripk3* rather than *Caspase-8* nearly completely corrected immunodeficiency disorders, developmental defects, and necroptosis in the thymus of RIPK1-deficient rats. However, inflammatory enteritis was still detected in either *Ripk1*/*Ripk3*-or *Ripk1*/*Casp-8*-double deficient rats and only rescued in *Ripk1*/*Ripk3*/*Caspase-8*-triple deficient rats, suggesting RIPK3 mediated necroptosis and Caspase-8 dependent apoptosis contribute in different part in RIPK1-deficient caused the syndrome. Moreover, RIPK1-deficient rat dermal fibroblasts (RDFs) showed sensitive necroptosis and impaired NF-κB activation, also reported in *RIPK1-deficient* human patients. And pharmacological inhibition of NF-κB activation could sensitize necroptosis of rat cells. Since mutations that damage NF-κB activation could cause immunodeficiency in human patients, it suggested that aberrantly activated necroptosis play a vital role in certain kinds of primary immunodeficiency syndromes and pharmacological inhibition of necroptosis could be a novel therapeutic strategy for those diseases.

## Introduction

RIPK1 is a critical regulator of programmed cell death and inflammation downstream of death receptor super-families^1-4^, toll-like receptors (TLRs^5-7^), and interferon receptors activation^8-11^. The activated receptors engaged RIPK1 and sequentially formed various RIPK1-containing signaling complexes, which controlled RIPK1 function by varied post-translational modifications^8^. For example, in tumor necrosis factor (TNF) signaling, RIPK1 was recruited to membrane-bound TNFR1 signaling complex-I and other signaling molecules such as TRADD, TRAF2, and cIAP1/2. In complex-I, RIP1 was modified by cIAP1/2 with non-degradative polyubiquitylation, which allows recruitment and activation of TAK1 to induce canonical NF-κB and MAP kinase signaling pathways. It leads to cell proliferation and inflammation, independent of RIPK1 kinase activity^3^. When the polyubiquitylation of RIPK1 was removed by the deubiquitinase cylindromatosis (CYLD) and A20, RIPK1 was engaged in a soluble signaling complex-II containing FADD, Caspase-8, and RIPK3^12^. Then RIPK1 kinase is activated to promote RIPK1-RIPK3 interaction and further aggregation to form a sizeable ∼2MDa complex (referred to as ripoptosome) which leads to cell apoptosis or necroptosis^3,13-15^.

*Ripk1* knockout mice died within postnatal day 3 with systemic inflammation and fatal respiratory failure^1,16,17^, suggesting the essential function of RIPK1 in development. While recently, homozygous loss-of-function mutations in the human RIPK1 have been described to cause immunodeficiency and intestinal inflammation in children ^18,19^. Cells from these child patients have impaired response to TNF to induce canonical NF-κB activation, suggesting the deficiency of the NF-κB pathway is probably the cause of the immunodeficiency symptoms. Consistently, impairment of upstream regulators of the NF-κB path, such as NF-κB essential modulator (NEMO) or the component of the linear ubiquitination chain assembly complex (LUBAC), which induces NEMO linear ubiquitination, or gain of function mutations on the inhibitor of NF-kB alpha (IκBα) will cause combined autoimmunity and immunodeficiency symptoms in human ^20-23^. In contrast to humans, disrupting the X-linked gene encoding NEMO in mice produces male embryonic lethality ^24^. Deficiency of mouse RIPK1 or LUBAC components will also cause perinatal or embryonic lethality ^16,17,25^. It suggests NF-κB signaling is essential for survival in mice but not in humans, which may originate in the genetic variation between inbred mouse strains and the highly heterogeneous human population. So, studies on the physiological functions of RIPK1 or other regulators of the NF-kB pathway in outbred animal models will give more insights into NF-κB dysfunction caused by autoimmunity and primary immunodeficiency.

In this study, we generated the RIPK1-deficient rat. These animals could reproduce the pathological phenotypes of the relative human child patients as inflammatory enteritis combined with immunodeficiency, characterized by T cell lymphopenia with abnormal thymus morphology and necroptosis in the thymus. These rats died during the late neonatal period. Compared to wild-type fibroblasts, the RIPK1-deficient rat fibroblasts show impaired NF-κB signaling. Interestingly, rats with combined *Ripk1* and *Ripk3* deficiency displayed normal thymotic development, and both T cell lymphopenia and necroptosis in the thymus were rescued. While fibroblasts derived from these rats still had impaired NF-κB signaling, suggesting RIPK3-dependent necroptosis but not the dysfunction of NF-κB activated gene-transcription is the critical cause of RIPK1-deficient induced immune deficiency, unlike *Ripk1/Casp-8* double knockout rats, which also exhibited late neonatal lethality, most *Ripk1/Ripk3* double knockout rats could survive to adulthood. All these RIPK1/RIPK 3 deficient rats still suffered from inflammatory enteritis. While the intestine is normal in mice with combined knockout of *Ripk1, Ripk3*, and *Casp-8*, suggesting RIPK1-deficiency-induced inflammatory enteritis was caused by both RIPK3-induced necroptosis and Caspase-8-induced apoptosis. Besides RIPK1 deficiency, pharmacological inhibition or genetic deletion upstream activators of NF-κB will accelerate necroptosis signaling and cause human immunodeficiency syndromes ^26-29^. Therefore, our rat model suggested that RIPK3-dependent necroptosis is the primary cause of RIPK1-deficient induced immunodeficiency and may contribute to other NF-κB dysfunction-associated immunodeficiency. Necroptosis will be the therapeutic target for treating these primary immune deficiency diseases (PIDs).

## Results

### RIPK1-deficient rats suffered from lymphopenia

To elucidate the physiological function of RIPK1 in outbred animals, we generated RIPK1-deficient rats (Fig. S1a). The RIPK1-deficient rats were progressively weakened after postnatal day three. They showed growth failure compared with wild-type littermates, which had also been reported in 8 PIDD child patients caused by RIP1 homozygous mutations (Fig.1 a and b) ^30^. Most RIPK1-deficient rats died during the late neonatal period (Fig. S1b). Cell counting showed that the *Ripk1*^-/-^ rats displayed T cell lymphopenia (Fig.1c and S1c), a common symptom of RIPK1-deficient child patients^18,30^. Besides that, these rats showed an increment of neutrophils which suggested suffering from infection or noninfectious inflammation (Fig. 1d). Similar results of blood assays were also reported in the cases of child patients^18^. We also checked the lymphoid organs, such as the spleen and thymus. Both of thymus and spleen of *Ripk1*^-/-^ rats were smaller than that of wild-type littermates (Fig.1e). While only the thymus-weight-to-body-weight ratio but not the spleen-weight-to-body-weight ratio of *Ripk1*^-/-^ rats were significantly smaller than wild-type littermates (Fig.1 f and g), indicating the development of thymus was abnormal (Fig.1 e to g). Interestingly, smaller thymus has been reported as a common phenomenon in a classical immunodeficient animal model such as SCID (server combined immune-deficiency) mice^31^, which suggests RIPK1-deficient will also cause a developmental defect in the thymus. And it was confirmed by the fact that the total thymocytes of *Ripk1*^-/-^ rats were much rare compared with *Ripk1*^+/-^ littermates (Fig. 1h). Consistently, the hematoxylin-eosin (H&E) staining of rat thymus, which showed muddy boundary of thymic cortex and thymic medulla with abnormally decreased thymic cortex area in Ripk1^-/-^ rats (Fig. 1i). And T cell development in the thymus was also abnormal in RIPK1-deficient rats (Fig. 1j, and Fig. S1 d and e). It is worth noting that the proportion of CD4^+^CD8^+^ double positive (DP) T cells decreased, and CD4^+^CD8^-^ or CD4^-^ CD8^+^ single positive (SP) T cells increased in RIPK1-deficient rats, suggesting defects occurred at the stage of T cells differentiation from DP cells to SP cells (Fig. S1 d and e). Consisted with that, apoptotic signal marked by cleaved Caspase-3 dramatically increased in the thymic cortex of RIPK1-deficient rats, suggesting injury in these regions caused excessive DP T cells to die, which showed abnormally differentiation (Fig. 1k). Therefore, these developmental defects in thymus probably were the direct cause of *Ripk1*-deficiency induced T cell lymphopenia.

**Figure 1:**
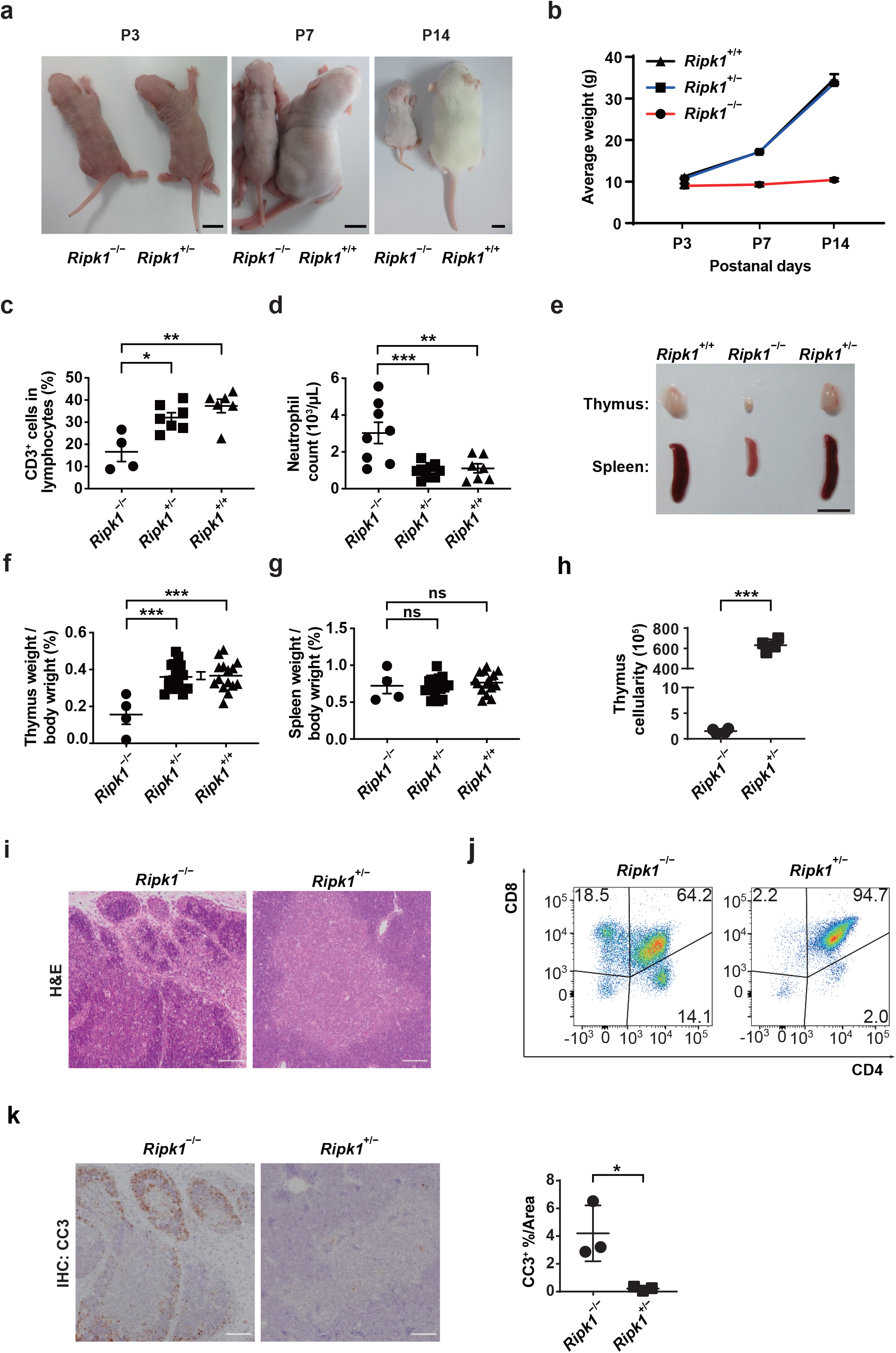
RIPK1-deficient neonatal rats suffered from immunodeficiency combined with inflammatory symptoms. (a) Representative images of a *Ripk1*^-/-^ rat and a *Ripk1*^+/-^ or *Ripk1*^+/+^ littermate at postnatal day 3, 7 and 14 (P3, P7 and P14) (scale bar = 1 cm). (b) Growth curve of body weight of *Ripk1*^-/-^ rats (n = 3 at P3, n=5 at P7, n=3 at P14) and *Ripk1*^+/-^ (n = 7 at P3, n=19 at P7, n=11 at P14) or *Ripk1*^+/+^ (n = 7 at P3, n=18 at P7, n=8 at P14) littermates. (c) Flow cytometric analysis of blood cells stained with anti-CD3 taken from P7 rats of the indicated genotypes. CD3^+^ cell proportion from *Ripk1*^-/-^ rats (n =4) was compared with *Ripk1*^+/-^ (n = 7) or *Ripk1*^*+/+*^ (n = 6) rats. (d) Neutrophil cells quantified using the ProCyte Dx hematology analyzer. Neutrophil number in whole blood from *Ripk1*^-/-^ rats (n = 8) was compared with *Ripk1*^+/-^ (n = 10) or *Ripk1*^+/+^ (n = 7) rats at P7. (e) Representative images of the thymus and spleen of a *Ripk1*^-/-^ rat or littermate controls at P7. Scale bar, 1 cm. (f and g) the weight of thymus (f) or spleen (g) relative to body weight of *Ripk1*^-/-^ rats (n = 4) were compared with *Ripk1*^+/-^ (n = 16) or *Ripk1*^+/+^ (n = 15) littermates at P7. (h) The absolute thymus cellularity of *Ripk1*^-/-^ rats or *Ripk1*^+/-^ littermates at P7. (n = 4, for each group). (i) Representative images of H&E staining on thymus sections from *Ripk1*^-/-^rats or *Ripk1*^+/-^ littermates at P7. Scale bar, 50 µm. (j) One example of flow cytometric analysis of thymocytes stained with anti-CD4 and anti-CD8 taken from *Ripk1*^-/-^rats or *Ripk1*^+/-^ littermates at P7. The statistical results were shown in supplementary Fig S1d and S1e. (k) Representative images of IHC staining with anti-cleaved-caspase-3 (CC3) antibody on thymus sections from *Ripk1*^-/-^ rats or *Ripk1*^+/-^ littermates at P7. Scale bar, 50 µm. Corresponding quantifications of the CC3 positive (brown) area on sections are shown on the right. Each dot represents the data from an individual rat (n = 3; for each group). Five pictures of every rat were selected and calculated by image J. All error bars and S.E.M. *P*-values were determined using the way-ANOVA (c, d, f, g) or Student’s *t-* test (h, k). *, *P* < 0.05; **, *P* < 0.01, ***, *P* < 0.001.

### RIPK1-deficient rats show inflammatory injury in multiple organs, including the intestine

Besides immunodeficiency-associated symptoms, RIPK1-deficient child patients were reported to suffer from early onset inflammatory bowel disease with different severities^18,30^. Simultaneously, RIPK1-deficient rats also develop enteritis characterized by epithelial cell death and abnormal villus structure (Fig. 2a). Besides that, RIPK1-deficient rats had epidermal hyperplasia with aberrant expression of keratin-14, which is confined to hair follicles in healthy skin (Fig. 2b)^17^. And lung injury showed with pulmonary hemorrhage, and diffuse alveolar damage was found in RIPK1-deficient rats during the late neonatal period, especially in the dying RIPK1-deficient rats, suggesting dyspnea led to their late-neonatal death (Fig. 2c). While in the early neonatal period, RIPK1-deficient rats showed simplified large alveoli and reduced area available for gaseous exchange in lung suggested by longer mean linear intercept lengths (Lm), which is similar as experimental Bronchopulmonary Dysplasia(BPD) rats (Fig. 2d) ^32^. Interestingly, liver injury was not apparent in RIPK1-deficient rats, which was quite different from the RIPK1-deficient mice (Fig. S2a)^17^.

**Figure 2:**
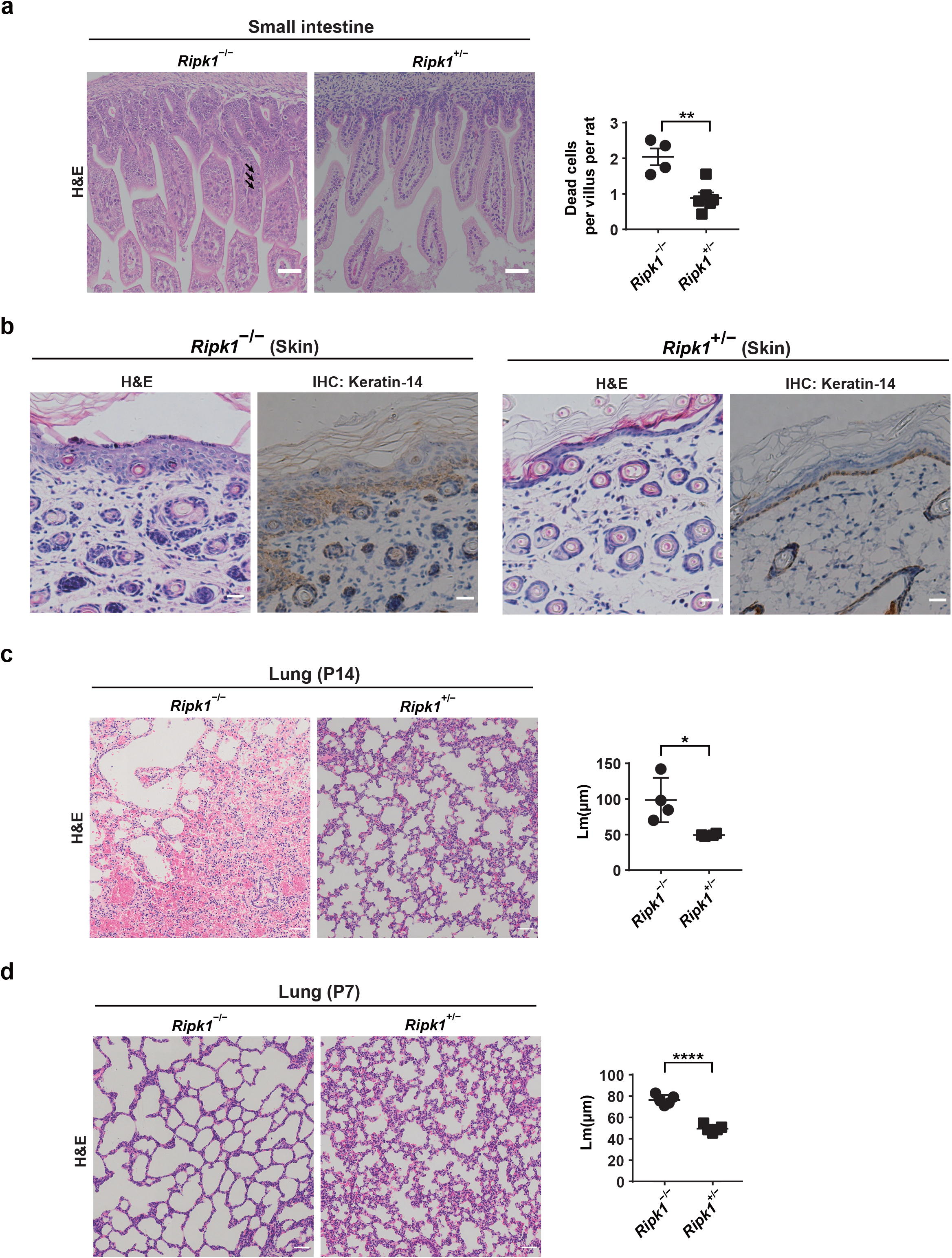
RIPK1-deficient neonatal rats showed multiorgan necroptosis. (a) Representative images of H&E staining on small intestine sections from *Ripk1*^-/-^ rats or *Ripk1*^+/-^ littermates at P7. Scale bar, 50 µm. Corresponding quantifications of dead cells (black arrow) per villus on sections were shown on the right. Each dot represented the data from an individual rat (*Ripk1*^-/-^ rats, n = 4 and *Ripk1*^+/-^ rats, n = 6). Five Pictures of every rat were selected, and more than 30 villi were calculated by Image J. (b) Representative images of H&E staining and IHC staining with anti-keratin14 antibody (brown) on skin sections from Ripk1^-/-^ rats or Ripk1^+/-^ littermates at P14. Scale bar, 50 µm. (c and d) Representative images of H&E staining on lung sections from *Ripk1*^-/-^ rats or *Ripk1*^+/-^ littermates at P14 (c) or P7(d). Scale bar, 50 µm. Corresponding quantifications of the mean linear intercept length (Lm), which represented the complexity of the air space in the lung on sections, were shown on the right. Each dot represented the data from an individual rat (n=4; for each group at P14, n=5; for each group at P7). Five pictures of every rat were selected, and Lm was calculated by image J. Error bars, and S.E.M. *P*-values were determined using Student’s *t-*test (a, c, d). *, *P* < 0.05; **, *P* < 0.01; ****, *P* < 0.0001;

### RIPK1-deficient caused lymphopenia can be resecured by knocking out *Ripk3*

According to a previous study, systemic inflammatory injury of RIPK1-deficient mice is mediated by RIPK3 and Caspase-8, which will lead to perinatal lethality ^16,17^. We then generated double knock-out of *Ripk1*/*Ripk3* or *Ripk1*/*Casp-8* in rats (Fig.S3 a and b) and found that only *Ripk*3 knock-out could extend the survival of RIPK1-deficient rats to adulthood (Fig.S3 c and d). These *Ripk1/Ripk3* double knock-out rats displayed a similar appearance as wild-type littermates (Fig.3 a and b). The increment of neutrophils and decrement of peripheral T cells were restored in young *Ripk1*^-/-^*Ripk3*^-/-^ rats, suggesting immunodeficiency combined with system inflammation were rescued by additionally knocking-out *Ripk3* (Fig. 3 c and d). Consistently, the thymus in young *Ripk1*^-/-^*Ripk3*^-/-^ rats was in normal development, and T cell differentiation in the thymus was also normal (Fig.3 e to g, and S3 e to f). What’s more, necroptosis signal (MLKL p-S345) is widespread in the thymus of baby *Ripk1*^-/-^ rats but scarce in young *Ripk1*^-/-^*Ripk3*^-/-^ rats, suggesting developmental abnormal in *Ripk1*^-/-^ rats thymus was probably due to necroptosis (Fig.3 h and i, S3 g and h). Besides the thymus, the injury in the lung was also recovered in *Ripk1*/*Ripk3* -deficient rats (Fig.S3i). Nevertheless, these rats are not healthy as wide-type rats. *Ripk1*^-/-^*Ripk3*^-/-^ young rats still suffer intestine epithelial cell death (Fig.3j). Interestingly, most dying *Ripk1*^-/-^*Ripk3*^-/-^ rats have persistent diarrhea. Severe inflammatory injury of enteritis was detected by histopathological analysis, suggesting enteritis may be the primary cause of death of old *Ripk1*/*Ripk3*-deficient rats (Fig. 4 a and b). Reduce expression of Caspase-8 in *Ripk1*^-/-^*Ripk3*^-/-^*Casp*-8^-/-^or *Ripk1*^-/-^ *Ripk3*^-/-^*Casp*-8^+/-^ rats could rescue enteritis and extend the survival rate of *Ripk1*^-/-^*Ripk3*^-/-^ rats, suggesting *Casp*8 mediated apoptosis contributed enteritis (Fig. 4 a and b). To explore if RIPK1 deficiency caused enteritis solely depends on *Casp8-mediated* apoptosis, we checked the intestine of *Ripk1*^-/-^*Casp*-8^-/-^ rats. It still showed enteritis characterized by a significant increment of dead epithelial cells and loss of villus structure compared with *Ripk1*^+/-^*Casp*-8^+/-^littermates (Fig. 4c). In addition, *Ripk1*^-/-^*Casp*-8^-/-^rats showed decreased peripheral T cells and increased neutrophils in blood (Fig.4 d and e), similar to RIPK1 deficient rats. It suggested T cell lymphopenia and system inflammation were not rescued by knocking-out *Casp-8*. So, these results distinguished the RIPK1-deficient caused symptoms as that associated immunodeficiency syndromes mainly relayed on RIPK3 dependent necroptosis and, enteritis injuries are both RIPK3 and CASP8 dependent.

**Figure 3:**
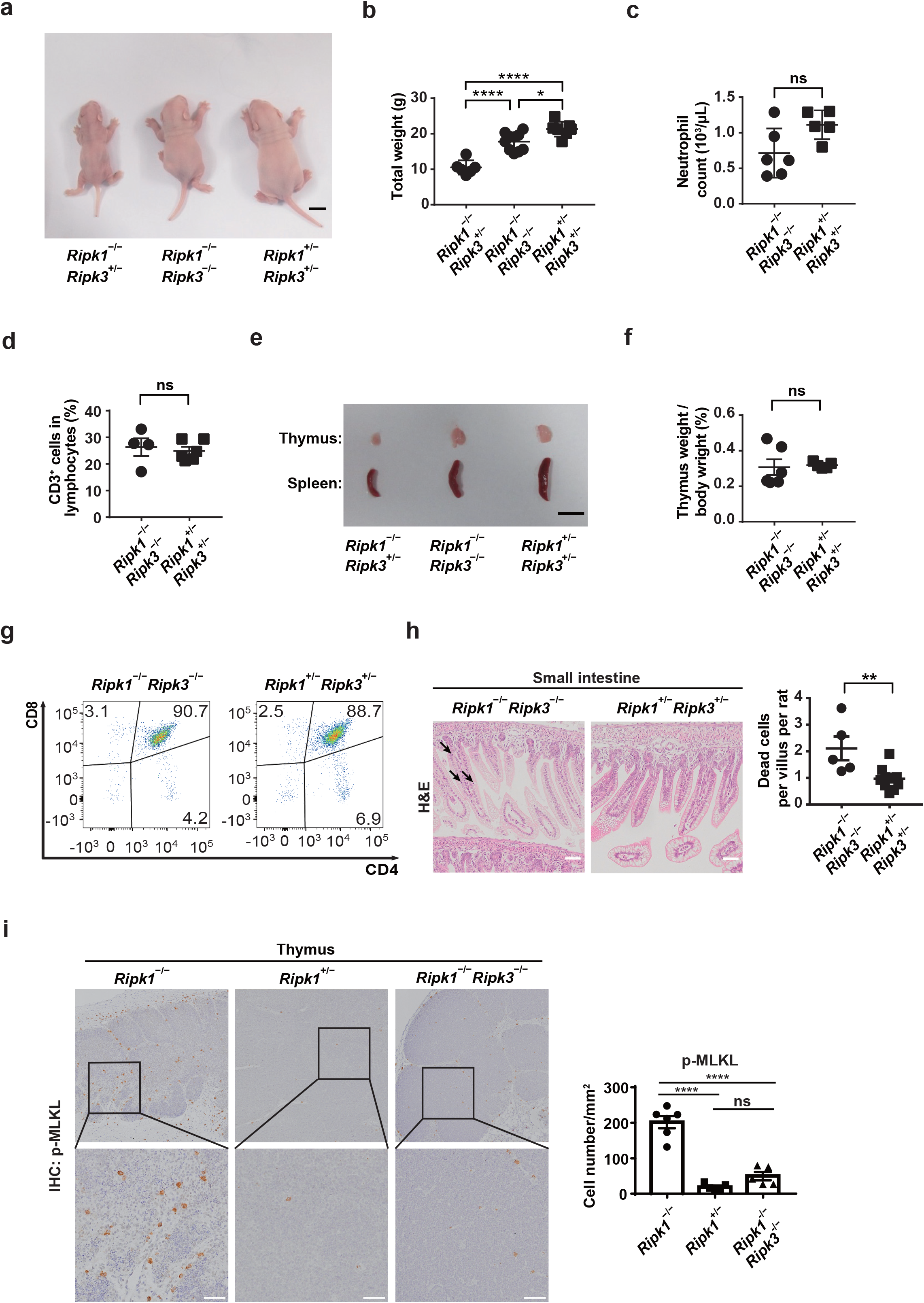
Knocking out *Ripk3* rescued RIPK1-deficient neonatal rats from inflammatory symptoms and early death. (a)Representative images of a *Ripk1*^-/-^*Ripk3*^+/-^, *Ripk1*^-/-^*Ripk3*^-/-^or *Ripk1*^+/-^ *Ripk3*^+/-^ littermate at P7 (scale bar = 1cm). (b) Body weight of indicated rats at P7. *Ripk1*^-/-^*Ripk3*^+/-^ rats (n=6) were compared with *Ripk1*^-/-^*Ripk3*^-/-^ (n=10) and *Ripk1*^+/-^*Ripk3*^+/-^ rats (n=7). (c) Neutrophil cells quantified using the ProCyte Dx hematology analyzer. Neutrophil number in whole blood from *Ripk1*^-/-^*Ripk3*^-/-^ rats (n = 6) was compared with *Ripk1*^+/-^*Ripk3*^+/-^ (n = 5) rats at P7. (d) Flow cytometric analysis of blood cells stained with anti-CD3 taken from P7 rats of the indicated genotypes. CD3^+^ cell proportion from *Ripk1*^-/-^*Ripk3*^-/-^ rats (n = 4) was compared with *Ripk1*^+/-^*Ripk3*^+/-^ (n = 6) rats at P7. (e) Representative images of the thymus and spleen of a *Ripk1*^-/-^*Ripk3*^-/-^ rat or littermate controls at P7 (scale bar = 1cm). (f) The weight of thymus relative to body weight of *Ripk1*^-/-^*Ripk3*^-/-^ rats (n = 6) were compared with *Ripk1*^+/-^*Ripk3*^+/-^ littermates (n=5) at P7. (g) One example of flow cytometric analysis of thymocytes stained with anti-CD4 and anti-CD8 taken from *Ripk1*^-/-^ *Ripk3*^-/-^ rats or *Ripk1*^+/-^*Ripk3*^+/-^ littermates at P7. The statistical results were shown in supplementary Fig S3f and S3g. (h) Representative images of H&E staining on small intestine sections from *Ripk1*^-/-^*Ripk3*^-/-^ rats or *Ripk1*^+/-^*Ripk3*^+/-^ littermates at P7. Scale bar, 50 µm. Corresponding quantifications of dead cells (black arrow) per villus on sections were shown on the right. Each dot represented the data from an individual rat (*Ripk1*^-/-^ *Ripk3*^-/-^ rats, n=5; *Ripk1*^+/-^*Ripk3*^+/-^, n=10). Five Pictures of every rat were selected, and more than 30 villi were calculated by Image J. (i) Representative images of IHC staining with Anti-MLKL (phospho S345) antibody (*p*-MLKL) antibody on thymus sections from *Ripk1*^-/-^ rats or indicated littermates at P7: scale bar, 50 µm. Corresponding quantifications of *p*-MLKL positive cells (brown) from Fig 3i were shown on the right. Each dot represents the data from an individual rat (*Ripk1*^-/-^ rats, n=6; *Ripk1*^+/-^ rats, n=5 and *Ripk1*^-/-^ *Ripk3*^-/-^ rats, n=5). Five pictures of every rat were selected and calculated by image J. (i) All error bars, S.E.M. *P*-values were determined using the one way-ANOVA (b, i) or Student’s *t-*test (c-d, f, h). *, *P* < 0.05; **, *P* < 0.01, ****, *P* < 0.0001.

**Figure 4:**
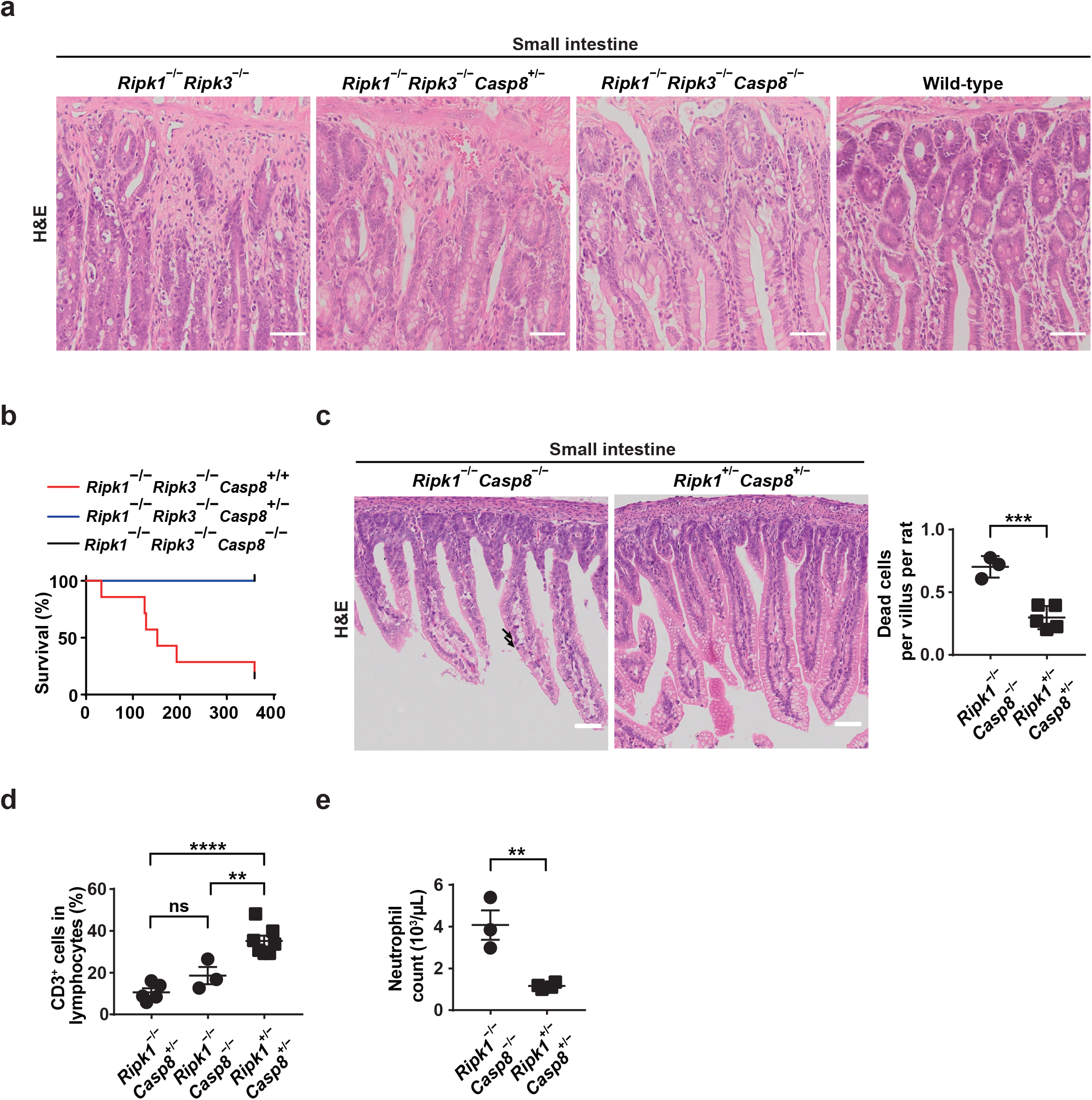
Caspase 8 knockout could rescue inflammation in *Ripk1*^-/-^*Ripk3*^-/-^ rats. (a) Representative images of H&E staining on small intestine sections from *Ripk1*^-/-^*Ripk3*^-/-^ rats or indicated littermates at 20 months. Scale bar, 50 µm. (b) Kaplan-Meier survival curve of indicated rats after birth. *Ripk1*^-/-^ *Ripk3*^-/-^*Casp8*^+/+^ rats (n = 14, red) were compared with *Ripk1*^-/-^ *Ripk3*^-/-^*Casp8*^+/-^ rats (n = 5, blue) and *Ripk1*^-/-^ *Ripk3*^-/-^*Casp8*^-/-^rats (n = 6, black). (c) Representative images of H&E staining on small intestine sections from *Ripk1*^-/-^*Casp8*^-/-^ rats or *Ripk1*^+/-^ *Casp8*^*+*/-^ littermates at P7. Scale bar, 50 µm. Corresponding quantifications of dead cells (black arrow) per villus on sections were shown on the right. Each dot represented the data from an individual rat (*Ripk1*^-/-^*Casp8*^-/-^ rats, n=3 and *Ripk1*^+/-^ *Casp8+*^/-^ rats, n=5). Five Pictures of every rat were selected. More than 30 villi were randomly selected in each rat using Image J. (d) Flow cytometric analysis of blood cells stained with anti-CD3 taken from P7 rats of the indicated genotypes. CD3^+^ cell proportion from *Ripk1*^-/-^ *Casp8*^+/-^ rats (n =5) was compared with *Ripk1*^-/-^ *Casp8*^-/-^ rats (n = 3) and *Ripk1*^+/-^*Casp8*^+/-^rats (n = 7) rats. (e) Neutrophil cells quantified using the ProCyte Dx hematology analyzer. Neutrophil number in whole blood from *Ripk1*^-/-^ *Casp8*^-/-^ rats (n = 3) was compared with *Ripk1*^+/-^*Casp8*^+/-^ rats (n = 4) rats at P7. All error bars and S.E.M. *P*-values were determined using Student’s *t-*test (c, e) or one-way ANOVA (d). Error bars are average ±S.E.M. ns, not significant; **, *P* < 0.01; ***, *P* < 0.001; ****, *P* < 0.0001.

### NF-κB dysfunction is not the direct cause of immunodeficiency-associated symptoms in RIPK1-deficient rats

Our data showed immunodeficiency-associated symptoms in RIPK1-deficient rats could be rescued by knocking out of *Ripk3*. While previous reports showed that patients with disrupting expression of RIPK1 showed impaired NF-κB activation^18^, and kinds of mutations to disrupt NF-κB activation could also cause immunodeficiency symptoms in humans^20-22^. It may indicate that dysfunction of NF-κB immediately causes immunodeficiency in RIPK1-deficient rats. So, we analyzed NF-κB activation in rat dermal fibroblasts (RDF) stimulated with TNF-α. It showed impaired NF-κB activation in *Ripk1* deficient rats, consistent with studies on human and mouse cells (Fig. 5a and S4a)^1,18^. Interestingly, RDF from *Ripk1*^-/-^*Ripk3*^-/-^ rats also showed impaired NF-Κb activation (Fig. 5a and S4a). It indicated NF-κB dysfunction in *Ripk1*^-/-^*Ripk3*^-/-^ rats did not cause immunodeficiency-associated syndromes. So, in RIPK1-deficient rats, NF-κB dysfunction may not be the leading cause of immunodeficiency. The abnormal thymus and T cell lymphopenia, the main immunodeficiency symptoms in RIPK1-deficient rats, could be rescued by knocking out of *Ripk3*. And necroptosis, the core function of RIPK3, was abnormally enriched in the thymus of RIPK1-deficient rats but scarce in the *Ripk1*^-/-^ *Ripk3*^-/-^ or control *Ripk1*^+/-^ rats, suggesting necroptosis significant major role in the thymus injury and lymphopenia. Various inflammatory cytokines, including TNF or IFN family members, could also induce RIPK3-dependent necroptosis^17,33^. To clarify which cytokine intrinsically caused necroptosis spontaneous activation in RIPK1-deficient rats, we analyzed a bulk of circulating inflammatory cytokines in RIPK1-deficient rat serum under normal conditions (without stimulation). It showed that IFN-γ was abnormally increased in the blood of RIPK1-deficient rats; other cytokines such as IL-2, IL-6, and IL-12p70 were all at typically low levels or could not be detected in standard blood samples from *Ripk1*^-/-^ or *Ripk1*^+/-^ rat littermates (Fig. 5b and S4 b-d). Interestingly, necroptosis could be induced by treatment of IFN-γ to RDF cells from RIPK1-deficient rats, suggesting abnormally elevated IFN-γ may be one of the inducers to cause necroptosis in *Ripk1*^-/-^ rats (Fig. S4e).

**Figure 5:**
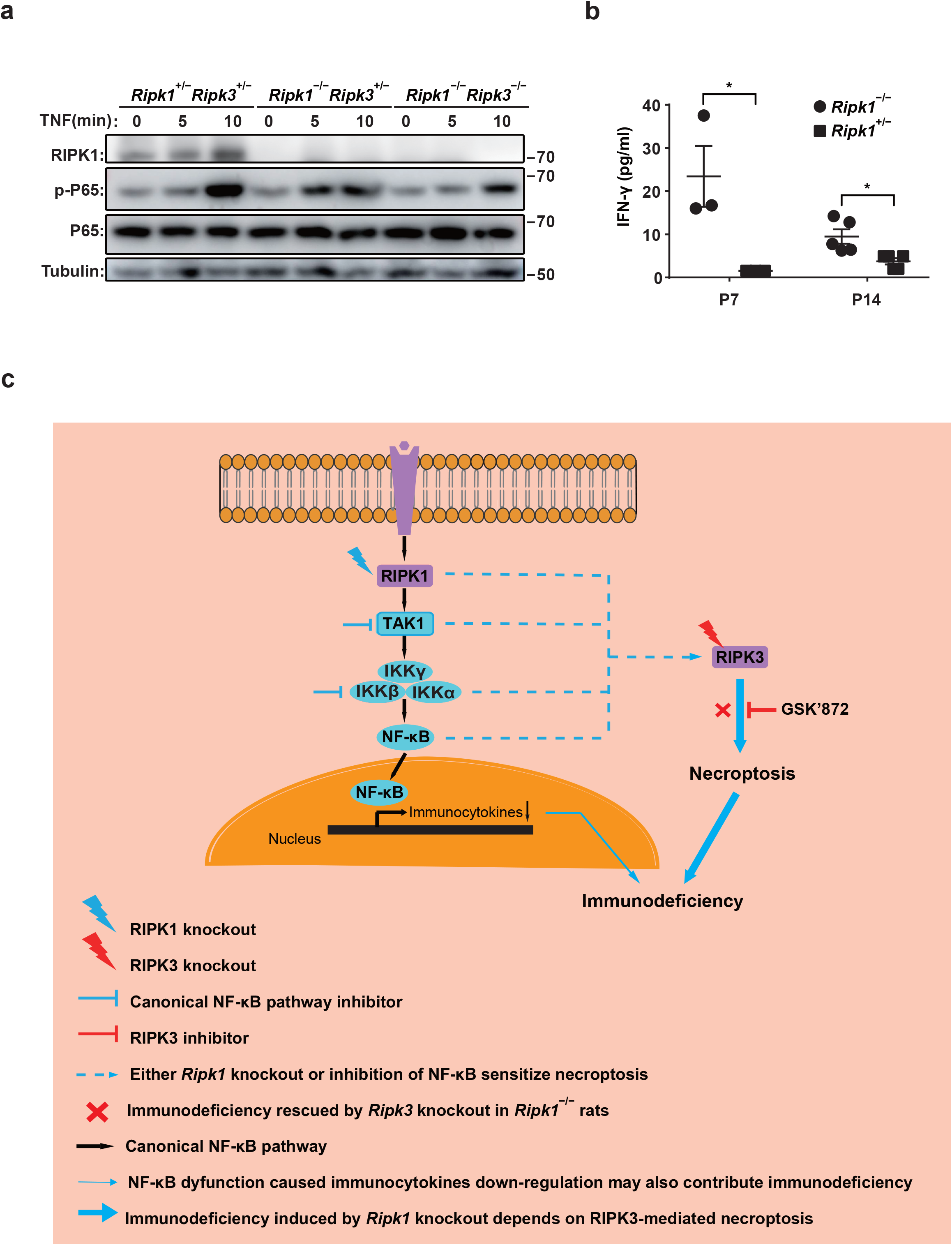
RIPK1-deficient caused NF-κB dysfunction was not rescued by *Ripk3* knocking-out. (a) Rat dermal fibroblasts (RDFs) from indicated rats were stimulated with 50 ng/mL human TNF-α and cell extracts were subjected to immunoblotting probed with antibodies that recognize a phosphorylated form of NF-κB (*p*-P65), NF-κB (P65), RIPK1, and β-tubulin. (b) Blood IFN-γ was assayed in untreated serums from rats at P7 (n=3; for each group) and P14 (n=5; for each group) by Liminux. (c) Schema graph shows that necroptosis is indispensable in the progression of immunodeficiency syndromes in RIPK1-deficient rats. In the canonical NF-κb pathway (black arrow), stimulus from the cell membrane will recruit RIPK1 and TAK1, activate the IKKβ complex and induce NF-κB activation and cytokine expression. (Blue line) In rat RDFs, when the NF-κB pathway was blocked by TAK1 inhibitor, IKK-β inhibitor, or knocking out *Ripk1*, RIPK3 activation, and necroptosis were sensitized. Similarly, in RIPK1-deficient rats (blue lightning), impaired NF-κB activation was also accompanied by necroptosis and immunodeficiency. *Ripk1*^-/-^ *Ripk3*^-/-^ rats (red lightning) could almost totally rescue the immunodeficiency in RIPK1-deficient rats while keeping impaired NF-κB pathway. All error bars and S.E.M. *P*-values were determined using two-way ANOVA followed by multiple comparisons (b). *, *P* < 0.05

## Discussion

Human primary immune deficiency diseases (PIDDs) are a heterogeneous group of disorders that impair the immune system. Most PIDDs result from monogenic inherited defects affecting various immune processes, such as immune development, effector cell activities, and maintaining immune homeostasis. Defects in signal pathways such as NF-κB, which are involved in the regulatory process of the immune response, are one of the sub-types of PIDDs characterized by a combination with autoimmunity symptoms, e.g., early-onset inflammatory bowel disease ^34,35^. These PIDDs arose from mutations impairing upstream regulators of NF-κB, including RIPK1 ^18,20-22^. Both innate and adaptive immune responses are under the control of the NF-κB family of transcription factors^36,37^. So immunodeficiency caused by impaired canonic NF-κB activation was initially thought to come from the distraction of NF-κB mediated transcription^23^. Our rat model showed that knocking out of *Ripk3* could nearly completely recover lost-of-RIPK1-caused immunodeficiency syndromes, including maldevelopment of the thymus and T cell lymphopenia (Fig.3d-g). It suggested that necroptosis mediated by RIPK3 plays a vital role in RIPK-deficiency-caused immunodeficiency syndromes. Consistently, the Mlkl p-S345, the marker of necroptosis activation, is intensive in the IHC of RIPK1-deficient rat thymus, while scarce in IHC of the control RIPK1^+/-^ rats (Fig. 3h and i). And cells from *Ripk1/Ripk3* double knockout rats showed impaired NF-κB activation comparable with *Ripk1* knockout cells (Fig. 4a and i). It indicated RIPK3-dependent necroptosis but not NF-κB dysfunction altered gene expression contributed directly to RIPK1-deficient induced immunodeficiency. Moreover, impairing NF-κB activation has been found to sensitize RIPK3-mediated necroptosis, suggesting the immune deficiency in patients with dysfunctional mutations in NF-κB-related genes may also be directly affected by abnormal necroptosis activation (Fig. S4f) ^26-29^. Therefore, it is imperative to verify if knocking out of Ripk3 could also benefit primary immune deficiency animals with other NF-κB dysfunctional mutations in the future, which will suggest necroptosis-dependent PIDDs (ND-PIDDs) as a new subtype of primary immune deficiency diseases (Fig.5c). If actual, pharmacological inhibition of necroptosis as a targeted treatment of ND-PIDDs will also be theoretically feasible and worth to study.

## Acknowledgment

We are grateful for advice from Dr. Xingxu Huang for animal-related research and Dr. Yanfei Liu for cell-related research (ShanghaiTech). We thank the staff members of the Molecular and Cell Biology Core Facility of ShanghaiTech and Cell Biology Facility and Molecular Imaging Core Facility SIBCB for technical support, and the staff members of the Molecular Imaging at the National Facility for Protein Science in Shanghai, Zhangjiang Lab for providing paraffin embedding machine and practical assistance. This work was supported by the National Natural Science Foundation of China (32050187 to L.S., 31571427 to H.W.) and the scientific research start-up from the School of Life Science, ShanghaiTech University to H.W.

## Reference

1 Kelliher, M. A. et al. The death domain kinase RIP mediates the TNF-induced NF-kappaB signal. Immunity 8, 297–303 (1998).

2 Festjens, N., Vanden Berghe, T., Cornelis, S. & Vandenabeele, P. RIP1, a kinase on the crossroads of a cell’s decision to live or die. Cell Death Differ 14, 400–410, doi:10.1038/sj.cdd.4402085 (2007).

3 Vandenabeele, P., Declercq, W., Van Herreweghe, F. & xsVanden Berghe, T. The role of the kinases RIP1 and RIP3 in TNF-induced necrosis. Sci Signal 3, re4, doi:10.1126/scisignal.3115re4 (2010).

4 Biton, S. & Ashkenazi, A. NEMO and RIP1 control cell fate in response to extensive DNA damage via TNF-α feedforward signaling. Cell 145, 92–103, doi:10.1016/j.cell.2011.02.023 (2011).

5 Najjar, M. et al. RIPK1 and RIPK3 Kinases Promote Cell-Death-Independent Inflammation by Toll-like Receptor 4. Immunity 45, 46–59, doi:10.1016/j.immuni.2016.06.007 (2016).

6 Kaiser, W. J. et al. Toll-like receptor 3-mediated necrosis via TRIF, RIP3, and MLKL. J Biol Chem 288, 31268–31279, doi:10.1074/jbc.M113.462341 (2013).

7 Meylan, E. et al. RIP1 is an essential mediator of Toll-like receptor 3-induced NF-kappa B activation. Nat Immunol 5, 503–507, doi:10.1038/ni1061 (2004).

8 Ofengeim, D. & Yuan, J. Regulation of RIP1 kinase signaling at the crossroads of inflammation and cell death. Nat Rev Mol Cell Biol 14, 727–736, doi:10.1038/nrm3683 (2013).

9 Thapa, R. J. et al. Interferon-induced RIP1/RIP3-mediated necrosis requires PKR and is licensed by FADD and caspases. Proc Natl Acad Sci U S A 110, E3109–3118, doi:10.1073/pnas.1301218110 (2013).

10 Yang, D. et al. ZBP1 mediates interferon-induced necroptosis. Cellular & Molecular Immunology 17, 356–368, doi:10.1038/s41423-019-0237-x (2019).

11 Ingram, J. P. et al. ZBP1/DAI Drives RIPK3-Mediated Cell Death Induced by IFNs in the Absence of RIPK1. The Journal of Immunology 203, 1348–1355, doi:10.4049/jimmunol.1900216 (2019).

12 de Almagro, M. C., Goncharov, T., Newton, K. & Vucic, D. Cellular IAP proteins and LUBAC differentially regulate necrosome-associated RIP1 ubiquitination. Cell Death Dis 6, e1800, doi:10.1038/cddis.2015.158 (2015).

13 Feoktistova, M. et al. A20 Promotes Ripoptosome Formation and TNF-Induced Apoptosis via cIAPs Regulation and NIK Stabilization in Keratinocytes. Cells 9, doi:10.3390/cells9020351 (2020).

14 Hu, H. et al. RIP3-mediated necroptosis is regulated by inter-filament assembly of RIP homotypic interaction motif. Cell Death Differ 28, 251–266, doi:10.1038/s41418-020-0598-9 (2021).

15 Zhang, Y. et al. The MLKL kinase-like domain dimerization is an indispensable step of mammalian MLKL activation in necroptosis signaling. Cell Death & Disease 12, doi:10.1038/s41419-021-03859-6 (2021).

16 Dillon, C. P. et al. RIPK1 blocks early postnatal lethality mediated by caspase-8 and RIPK3. Cell 157, 1189–1202, doi:10.1016/j.cell.2014.04.018 (2014).

17 Rickard, J. A. et al. RIPK1 regulates RIPK3-MLKL-driven systemic inflammation and emergency hematopoiesis. Cell 157, 1175–1188, doi:10.1016/j.cell.2014.04.019 (2014).

18 Cuchet-Lourenco, D. et al. Biallelic RIPK1 mutations in humans cause severe immunodeficiency, arthritis, and intestinal inflammation. Science 361, 810–813, doi:10.1126/science.aar2641 (2018).

19 Tao, P. et al. A dominant autoinflammatory disease caused by non-cleavable variants of RIPK1. Nature 577, 109–114, doi:10.1038/s41586-019-1830-y (2020).

20 Zonana, J. et al. A novel X-linked disorder of immune deficiency and hypohidrotic ectodermal dysplasia is allelic to incontinentia pigmenti and due to mutations in IKK-gamma (NEMO). Am J Hum Genet 67, 1555–1562, doi:10.1086/316914 (2000).

21 Boisson, B. et al. Immunodeficiency, autoinflammation and amylopectinosis in humans with inherited HOIL-1 and LUBAC deficiency. Nat Immunol 13, 1178–1186, doi:10.1038/ni.2457 (2012).

22 Courtois, G. et al. A hypermorphic IkappaBalpha mutation is associated with autosomal dominant anhidrotic ectodermal dysplasia and T cell immunodeficiency. J Clin Invest 112, 1108–1115, doi:10.1172/JCI18714 (2003).

23 Pasparakis, M. & Kelliher, M. Connecting immune deficiency and inflammation. Science 361, 756–757, doi:10.1126/science.aau6962 (2018).

24 Makris, C. et al. Female mice heterozygous for IKK gamma/NEMO deficiencies develop a dermatopathy similar to the human X-linked disorder incontinentia pigmenti. Mol Cell 5, 969–979, doi:10.1016/s1097-2765(00)80262-2 (2000).

25 Peltzer, N. et al. LUBAC is essential for embryogenesis by preventing cell death and enabling haematopoiesis. Nature 557, 112–117, doi:10.1038/s41586-018-0064-8 (2018).

26 Yang, L. et al. TAK1 regulates endothelial cell necroptosis and tumor metastasis. Cell Death Differ 26, 1987–1997, doi:10.1038/s41418-018-0271-8 (2019)

27 Pescatore, A., Esposito, E., Draber, P., Walczak, H. & Ursini, M. V. NEMO regulates a cell death switch in TNF signaling by inhibiting recruitment of RIPK3 to the cell death-inducing complex II. Cell Death Dis 7, e2346, doi:10.1038/cddis.2016.245 (2016).

28 Lamothe, B., Lai, Y., Xie, M., Schneider, M. D. & Darnay, B. G. TAK1 is essential for osteoclast differentiation and is an important modulator of cell death by apoptosis and necroptosis. Mol Cell Biol 33, 582–595, doi:10.1128/mcb.01225-12 (2013).

29 McComb, S. et al. cIAP1 and cIAP2 limit macrophage necroptosis by inhibiting Rip1 and Rip3 activation. Cell Death Differ 19, 1791–1801, doi:10.1038/cdd.2012.59 (2012).

30 Li, Y. et al. Human RIPK1 deficiency causes combined immunodeficiency and inflammatory bowel diseases. Proc Natl Acad Sci U S A 116, 970–975, doi:10.1073/pnas.1813582116 (2019).

31 <A severe combined immunodeficiency mutation in the mouse.pdf>.

32 van Haaften, T. et al. Airway delivery of mesenchymal stem cells prevents arrested alveolar growth in neonatal lung injury in rats. Am J Respir Crit Care Med 180, 1131–1142, doi:10.1164/rccm.200902-0179OC (2009).

33 Roderick, J. E. et al. Hematopoietic RIPK1 deficiency results in bone marrow failure caused by apoptosis and RIPK3-mediated necroptosis. Proc Natl Acad Sci U S A 111, 14436–14441, doi:10.1073/pnas.1409389111 (2014).

34 Allenspach, E. & Torgerson, T. R. Autoimmunity and Primary Immunodeficiency Disorders. J Clin Immunol 36 Suppl 1, 57–67, doi:10.1007/s10875-016-0294-1 (2016).

35 Carneiro-Sampaio, M. & Coutinho, A. Early-onset autoimmune disease as a manifestation of primary immunodeficiency. Front Immunol 6, 185, doi:10.3389/fimmu.2015.00185 (2015).

36 Paul, S. & Schaefer, B. C. A new look at T cell receptor signaling to nuclear factor-kappa B. Trends in Immunology 34, 269–281, doi:10.1016/j.it.2013.02.002 (2013).

37 Hayden, M. S. & Ghosh, S. NF-kappaB in immunobiology. Cell Res 21, 223–244, doi:10.1038/cr.2011.13 (2011).

