## Supplemental figures and legends for "RIPK3-dependent necroptosis is the primary cause of RIPK1-deficient-induced immunodeficiency"

#### Supplementary figure legends

**Figure S1: *Ripk1$^{-/-}$* rat died early with decreased lymphocytes.**

(a) Schematic strategy of generating *Ripk1* knockout rats using the CRISPR/Cas9 system (upper panel). One example of PCR analysis of tail DNA of wild-type, heterozygous and knockout rats (Lower panel). (b) Kaplan-Meier survival curve of indicated rats after birth. *Ripk1$^{-/-}$* rats (n = 17, red) was compared with *Ripk1$^{+/-}$* (n = 42, blue) or WT (n = 39, black) rats. (c) Flow cytometric analysis showed the blood cells stained with anti-CD3 taken from P7 rats of the indicated genotypes. CD3$^{+}$ cell number from *Ripk1$^{-/-}$* rats (n = 4) was compared with *Ripk1$^{+/-}$* (n = 6) or *Ripk1$^{+/+}$* (n = 5) rats. (d and e) Percentage of double positive (CD4$^{+}$CD8$^{+}$ cells, DP) thymocytes (d) or single positive (CD4$^{+}$CD8$^{-}$ cells or CD4$^{-}$CD8$^{+}$, SP) thymocytes (e) in total thymocytes from *Ripk1$^{-/-}$* rats (n=5) or *Ripk1$^{+/-}$* littermates (n=4) at P7 were analyzed by flow cytometric (according to fig.1j). Each dot represents the data from an individual rat. All error bars, S.E.M. *P*-values, were determined using the way-ANOVA (c) or Student's t-test (d, e). *, $P < 0.05$; **, $P < 0.01$.

**Figure S2: *Ripk1$^{-/-}$* rats showed MLKL activation and no apparent liver injury.**

(a) Representative images of H&E staining on liver sections from *Ripk1$^{-/-}$* rats or *Ripk1$^{+/-}$* littermates at P7. Scale bar, 50 μm.

**Figure S3: RIPK3 but not Caspase 8 deficiency could rescue RIPK1-deficient-caused immunodeficiency in rats**

(a and b) The schematic strategy of generating *Ripk3* (a) or *Casp8* (b) knockout rats using the CRISPR/Cas9 system (upper panel). One example of PCR analysis of tail DNA of wild-type, heterozygous and knockout rats (Lower panel). (c) Kaplan-Meier survival curve of indicated rats after birth. *Ripk1$^{-/-}$Ripk3$^{+/-}$* rats (n = 12, red) were compared with *Ripk1$^{-/-}$Ripk3$^{-/-}$* (n = 19, blue) rats and *Ripk1$^{+/-}$Ripk3$^{+/-}$* (n = 15, black) rats. (d) Kaplan-Meier survival curve of RIPK1Casp8 rats after birth. *Ripk1$^{-/-}$ Casp8$^{+/-}$*rats (n = 8, red) were compared with *Ripk1$^{-/-}$Casp8$^{-/-}$* rats (n = 8, blue) and *Ripk1$^{+/-}$Casp8$^{+/-}$* rats (n = 12, black). (e and f) Percentage of double positive (CD4$^{+}$CD8$^{+}$ cells, DP) thymocytes (e) or single positive (CD4$^{+}$CD8$^{-}$ cells or CD4$^{-}$CD8$^{+}$, SP) thymocytes (f) in total thymocytes from *Ripk1$^{-/-}$Ripk3$^{-/-}$* rats (n = 4) or *Ripk1$^{+/-}$Ripk3$^{+/-}$* rats (n = 5) at P7 were analyzed by flow cytometric (according to fig.3g). Each dot represents the data from an individual rat. (g) HEK-293T cells were co-transfected with lentiviral vectors encoding Myc-tagged rMLKL(169-end) with vector, Flag-tagged rRIPK3 or lag-tagged rRIPK3(K51A) as indicated,24 hours later, cells were harvested, and the whole-cell lysates were subjected to immunoblotting probed with antibodies that recognize a phosphorylated form of MLKL (*p*-MLKL), Flag, Myc or β-actin. (h) Representative images of IHC staining with the anti-Phospho-mMLKL-S345 antibody on a thymus section from a Ripk1-deficient rat at P7. Scale bar,100μm. Five different views were shown in higher resolution (close-up view). (i) Representative images of H&E staining on lung sections from *Ripk1$^{-/-}$Ripk3$^{-/-}$* rats or *Ripk1$^{+/-}$Ripk3$^{+/-}$* littermates at P7. Scale bar, 50 μm. Corresponding quantifications of the mean linear intercept length (Lm), which represented the complexity of the air space in the lung on sections, were shown on the right. Each dot represented the data from an individual rat (*Ripk1*$^{-/-}$*Ripk3*$^{-/-}$ rats, n = 5 and *Ripk1*$^{+/-}$*Ripk3*$^{+/-}$ rats, n = 5). Five pictures of every rat were selected, and Lm was calculated by image J. Error bars, and S.E.M. *P*-values were determined using the Student's *t*-test (e-g). Ns, not significant.

**Figure S4: Inhibition of NF-κB sensitized RIPK3-mediated necroptosis.**

(a) A histogram of the western blot results in figure 5a. The relative *p*-P65 signals (*p*-P65/P65) at indicated times were compared from *Ripk1*$^{-/-}$ *Ripk3*$^{+/-}$ RDFs, *Ripk1*$^{-/-}$ *Ripk3*$^{-/-}$ RDFs and *Ripk1*$^{+/-}$ *Ripk3*$^{+/-}$ RDFs (n=3, for each group). (b-d) Blood IL-4 (b), IL-6 (c), and IL-12p70(d) were assayed in untreated serums from rats at P7 (n=3, for each group) and P14 (n=5, for each group) by Liminux. (e) Cell viability of primary RDFs isolated from indicated rats was determined by measuring cellular ATP levels as described in the methods. Z, Z-VAD-FMK; GSK'872, Ripk3 kinase inhibitor. Analysis of each sample was performed in triplicate. (f) Cell viability of primary RDFs isolated from WT rats was determined by measuring cellular ATP levels as described in the methods. T, TNF-α; Z, Z-VAD-FMK; TAKi, TAK1 inhibitor; TPCA-1, IKKβ inhibitor; GSK' 872, RIPK3 inhibitor. Analysis of each sample was performed in triplicate. All error bars, and S.E.M. *P*-values, were determined using two-way ANOVA followed by multiple comparisons (b-e) or one-way ANOVA (f). ns, not significant; *, $P < 0.05$.****, $P < 0.0001$.

### Figure S1

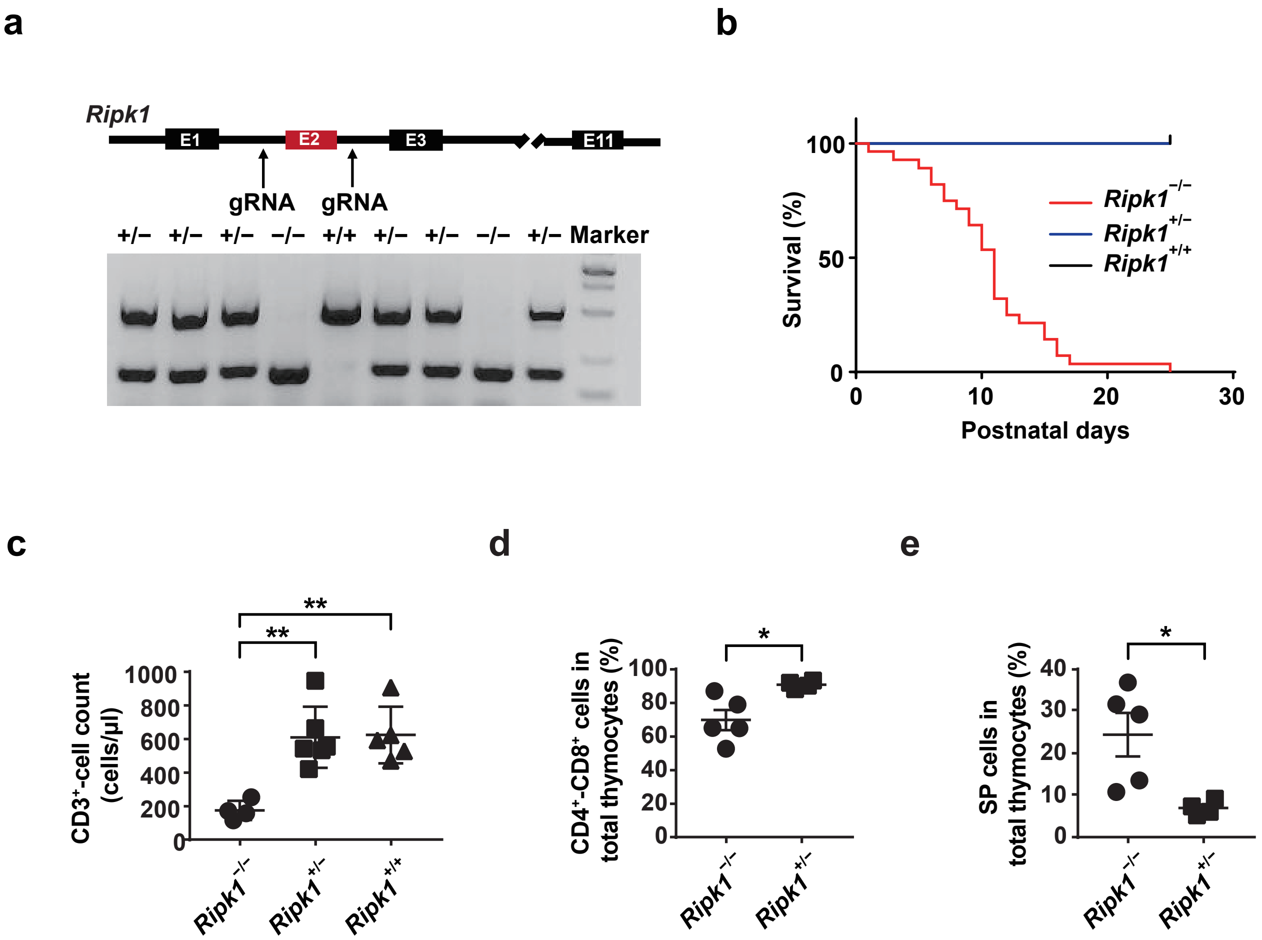
a
Ripk1
E1
E2
E3
E11
gRNA
gRNA
+/− +/− +/− −/− +/+ +/− +/− −/− +/− Marker
b
100
50
0
Survival (%)
0
10
20
30
Postnatal days
Ripk1−/−
Ripk1+/−
Ripk1+/+
c
CD3+-cell count (cells/µl)
1000
800
600
400
200
0
**
**
Ripk1−/−
Ripk1+/−
Ripk1+/+
d
CD4+-CD8+ cells in total thymocytes (%)
100
80
60
40
20
0
*
Ripk1−/−
Ripk1+/−
e
SP cells in total thymocytes (%)
40
30
20
10
0
*
Ripk1−/−
Ripk1+/−

**Figure S2**

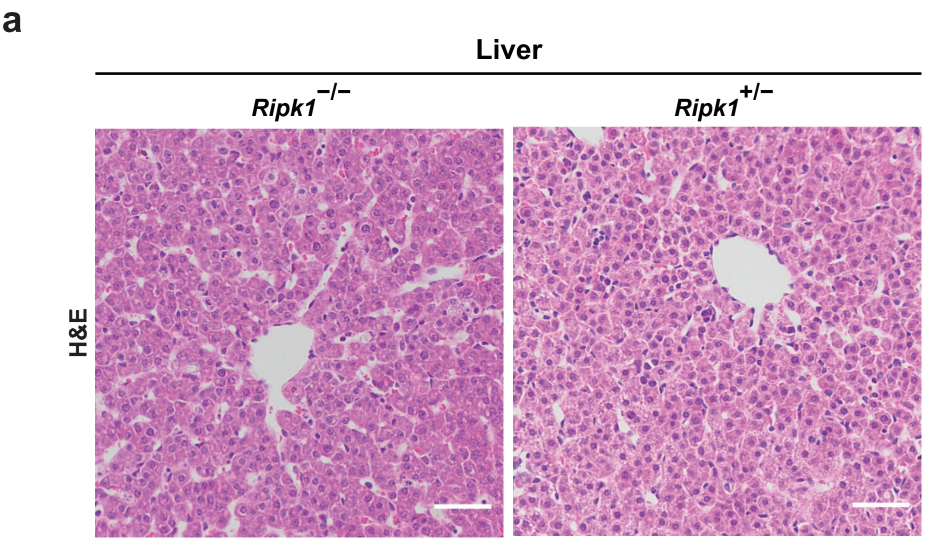

### Figure S3

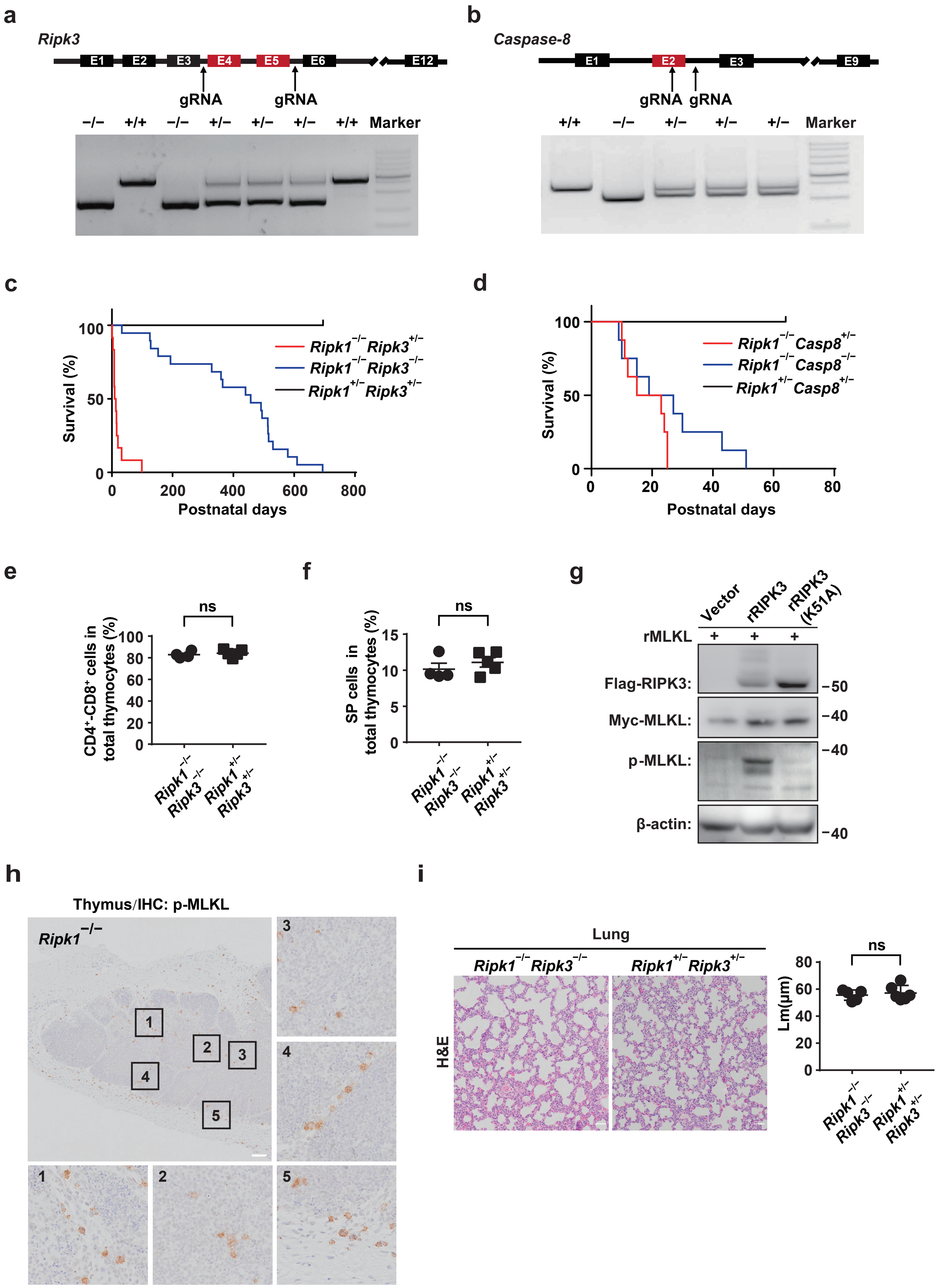

a
Ripk3
E1 E2 E3 E4 E5 E6 E12
gRNA gRNA
−/− +/+ −/− +/− +/− +/− +/+ Marker
b
Caspase-8
E1 E2 E3 E9
gRNA gRNA
+/+ −/− +/− +/− +/− Marker
c
Survival (%)
Postnatal days
Ripk1−/− Ripk3+/−
Ripk1−/− Ripk3−/−
Ripk1+/− Ripk3+/−
d
Survival (%)
Postnatal days
Ripk1−/− Casp8+/−
Ripk1−/− Casp8−/−
Ripk1+/− Casp8+/−
e
CD4+-CD8+ cells in total thymocytes (%)
ns
Ripk1−/− Ripk3−/−
Ripk1+/− Ripk3+/−
f
SP cells in total thymocytes (%)
ns
Ripk1−/− Ripk3−/−
Ripk1+/− Ripk3+/−
g
Vector
rRIPK3
rRIPK3 (K51A)
rMLKL + + +
Flag-RIPK3:
Myc-MLKL:
p-MLKL:
β-actin:
−50
−40
−40
−40
h
Thymus/IHC: p-MLKL
Ripk1−/−
1 2 3 4 5
i
Lung
Ripk1−/− Ripk3−/−
Ripk1+/− Ripk3+/−
H&E
Lm(μm)
ns

#### Figure S4

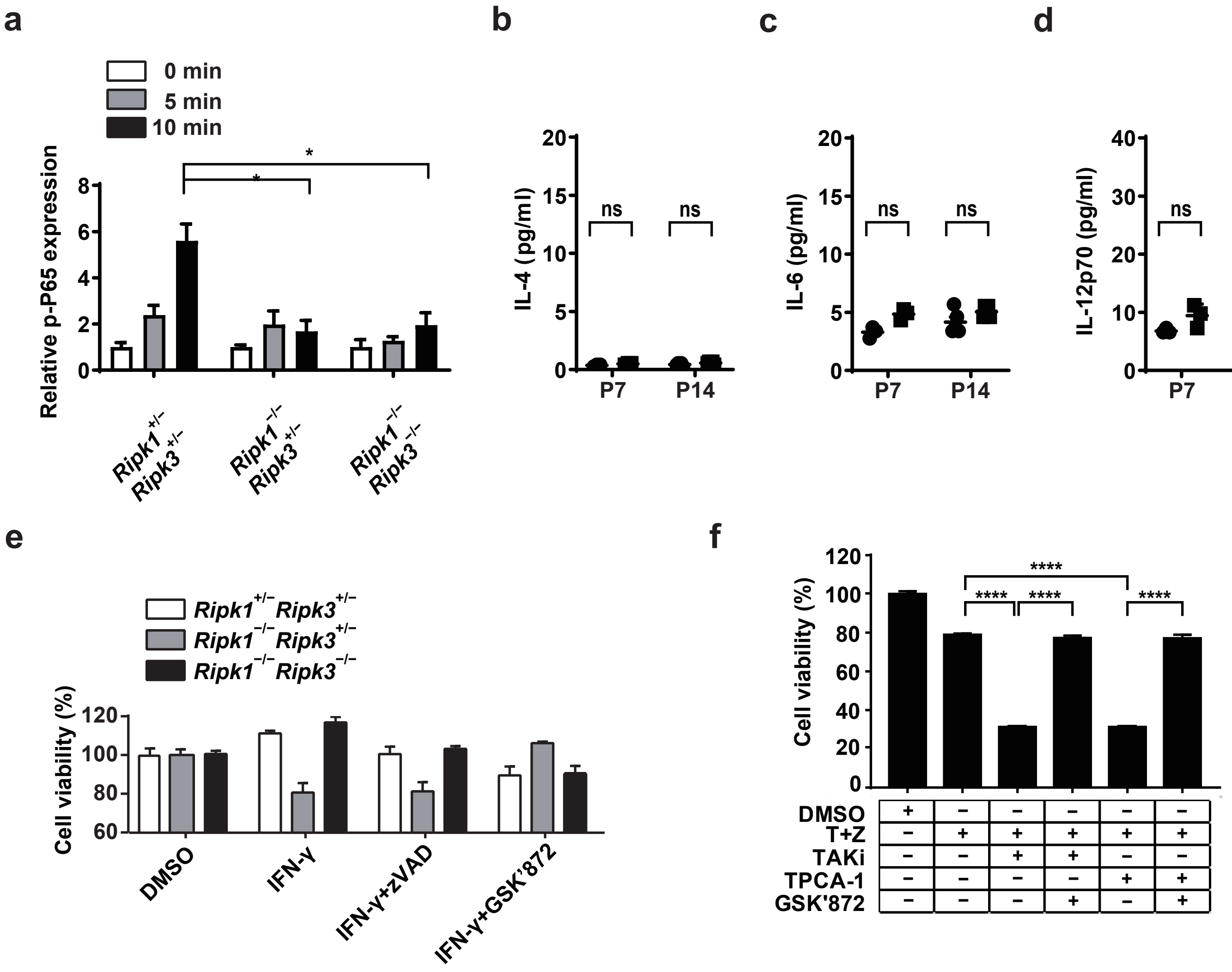

a
0 min
5 min
10 min
Relative p-P65 expression
*
*
Ripk1+/− Ripk3+/−
Ripk1−/− Ripk3+/−
Ripk1−/− Ripk3−/−
b
IL-4 (pg/ml)
ns
ns
P7
P14
c
IL-6 (pg/ml)
ns
ns
P7
P14
d
IL-12p70 (pg/ml)
ns
ns
P7
P14
e
Ripk1+/− Ripk3+/−
Ripk1−/− Ripk3+/−
Ripk1−/− Ripk3−/−
Cell viability (%)
DMSO
IFN-γ
IFN-γ+zVAD
IFN-γ+GSK'872
f
Cell viability (%)
****
****
****
****
DMSO
T+Z
TAKi
TPCA-1
GSK'872
